# Five seasons with DeepRootLab: A unique facility for easier deep root research in the field

**DOI:** 10.1101/2025.04.20.649645

**Authors:** Eusun Han, Corentin Clément, Weronika Czaban, Abraham George Smith, Dorte Bodin Dresbøll, Kristian Thorup-Kristensen

## Abstract

Deep-rooted crops accessing water and nutrients from deep soil layers enhance the resource base for crop production. However, studying these roots in field conditions is labor-intensive, limiting research scope. We established a field root research facility with 48 plots for replicated experiments. The facility includes 144 six-meter-long minirhizotron tubes and an AI-based pipeline for rapid root trait analysis. We also attempted to install access-tubes and customized ingrowth-core production for less-invasive root activity determination. Our study revealed significant differences in deep root density among species, particularly at depths of 2.5 to 4.5 meters, over five years. The less invasive studies using ingrowth-cores reached depths of 4.2 meters. Nutrient tracer 15N analysis showed marked differences in deep root activity among crop species. TDR sensors indicated varying water depletion in deeper soil layers, influenced by crop species and root growth patterns. We established a field facility for studying deep root growth and function, demonstrating its effectiveness in analyzing diverse deep-rooted plant species. This facility provides an ideal platform for conducting meaningful research in deep soil layers, yielding statistically and biologically significant results for agricultural applications.

## Introduction

No universal consensus has been reached on the pertinence of deep root development of crop plants for increased resource acquisition for food production (Thorup-Kristensen et al., 2020a). This has stemmed from the lack of detailed knowledge on the process of development and functioning of deep roots due to the difficulty in extending the research deeper into the arable subsoil under field conditions (Gregory et al., 2022). The requirements for labour, cost, and time to observe or sample the living roots in field conditions are high, therefore, often it acts as a bottleneck for deep root studies. Alternatively, semi-field (Van De Geijn *et al*., 1994; Svane *et al*., 2019b) and rhizobox-based facilities (Thorup-Kristensen *et al*., 2020b) have demonstrated the potential to study root systems for resource acquisition and root-soil interactions. Such facilities have the advantage of easier access and pre-installed equipment, such as water sensors, rhizotron tubes, or porous cups for soil water extraction. Further, they may allow a higher degree of control of soil and climatic factors, for example, drought conditions. While such approaches can offer more stable root measurements for detailed mechanisms at the root-soil interface, they do in many ways not fully represent real field conditions.

In field conditions, root researchers have to deal with natural soil with a high degree of physicochemical heterogeneity (Kautz *et al*., 2013). Deep installation of devices for measurements requires mechanical drilling with a high cost and is often restricted to a certain depth in the subsoil layers. The time-demanding process for root research *in situ* forces the research unit to minimize the range of treatments and temporal/spatial scales with a limited number of replicates. For instance, using the profile wall method, recording the rooting density up to 2 m of soil depth for 4-5 times during the season was possible only with two field plots per treatment (Huang *et al*., 2020; Han *et al*., 2021a). Similarly, soil sampling that requires arduous and time-consuming root extraction can also limit the number of temporal and spatial observation points. For example, the root-length density (RLD) measurement carried out from the soil cores was derived from two replicates only when root washing was involved (Kirkegaard *et al*., 2015). Four replicates were investigated when indirect measurement via root counting was carried out (Wasson *et al*., 2014). Such limitations caused by time-consuming sample collection can be solved with permanent structures for root measurement enabling non-destructive sampling.

It is challenging to acquire high-time resolution data on root growth. Destructive sampling may provide detailed answers at specific time points, but still relies on heavy labour for data collection. An alternative is the minirhizotron method (Rewald & Ephrath, 2013). It still requires the laborious installation of the minirhizotron tubes within each plot, but when this is done, it allows a rapid capture of root images. This leads to easier root observation in field conditions, for example, comparison of different crop species (Kristensen & Thorup-Kristensen, 2004) or cropping systems (Båth *et al*., 2008, p. 200) at multiple time points. A bottleneck of minirhizotrons has been extracting measurements from acquired images, which historically required labour-intensive grid-counting or manual annotation. It has been suggested that it takes approximately 20 minutes per meter of grid line (Böhm *et al*., 1977), which poses challenges for studies aiming at quantification of root growth at multiple observation points in time, space, and research treatments. Therefore, a more rapid root quantification method with high accuracy on field root images is necessary. Deep learning segmentation of root images has been proposed (Han *et al*., 2021b; Smith *et al*., 2022), which is yet to be validated in field conditions with a diverse range of species.

Crops have shown a wide range of deep rooting capacities. The major annual crops such as wheat and canola have been shown to establish root depths often to more than 2 m, depending on genotypes, phenology, and management practices (Thorup-Kristensen *et al*., 2009; Rasmussen *et al*., 2015; Han *et al*., 2015; Dresbøll *et al*., 2016). Perennial field crops such as lucerne, chicory and tall fescue are known to grow up to 2-3 m (Weaver, 1926; Han *et al*., 2021a). The emerging perennial grain crops such as intermediate wheatgrass and rosinweed are known to be deep-rooted below 2-3 m of depth (Van Tassel *et al*., 2017; Pugliese *et al*., 2019). However, continuous observation of deep-rooting perennial crops over a long period has not been reported sufficiently to understand species differences and their season-to-season as well as within-season dynamics. This information can be helpful when genotypic (G: genotype) potential for deep rooting and subsoil exploitation is in question, which is being attempted in semi-field conditions (Van De Geijn *et al*., 1994; Svane *et al*., 2019b). Choice of crops and time for sowing/harvest (M: management) greatly influence belowground production, and thus resource acquisition potential (Kristensen & Thorup-Kristensen, 2004; Rasmussen & Thorup-Kristensen, 2016). A high-resolution dataset in this regard has not been produced under field conditions. Therefore, such a research platform and facility are urgently called for GxM research (Hunt *et al*., 2021).

Therefore, we built a field facility, called **DeepRootLab**, that allows deep root studies to more than 4 m depth. Our aim in developing this facility was to generate statistically significant and biologically meaningful datasets with high efficiency. The previously published articles based on the data from DeepRootLab are shown in **Supp. Table 1**.

## Materials and methods

### Facility layout and crops

The DeepRootLab facility was established at the experimental station of the University of Copenhagen in Taastrup, Denmark (55 º 40’ N; 12 º 18’ E). The soil was classified as Agrudalf. A detailed description of the soil physical and chemical condition at the study site is available in Supp. Table 2. Weather conditions recorded at the study site are presented in Supp. Figure 1.

In total 24 experimental plots (one plot = 19.5 m x 10 m in size) were laid out in four blocks **(Figure 1)**. The 6 plots in each block were laid out in a 3 columns x 2 rows layout, and the plot heads of the two rows were facing each other at a distance of 4.5 m. The distance between neighbouring plots was 3 m. The distance between blocks was 5 m in both column and row-wise. We divided each plot into two kinds of plot layouts, namely intercropping setup (Layout I) and monocropping setup (Layout II). In layout I, 9 subplots were formed with alternating stripes, five 1.5 m-wide strips for perennial crops (1.5 m in width; 10 m in length) and four strips for annual crops next to each other (3 m in width; 10 m in length) for the study of crops in strip intercropping. In Layout II, the main plot was split into 2 subplots (9.75 m x 10 m), which can be used for individual crops or treatments. We chose and grew a wide range of perennial and annual crop species that are known to be deep-rooted as shown in **Table 1**. No data from intercropping treatments is presented here (see Supp. Table 1).

**Figure 1.**
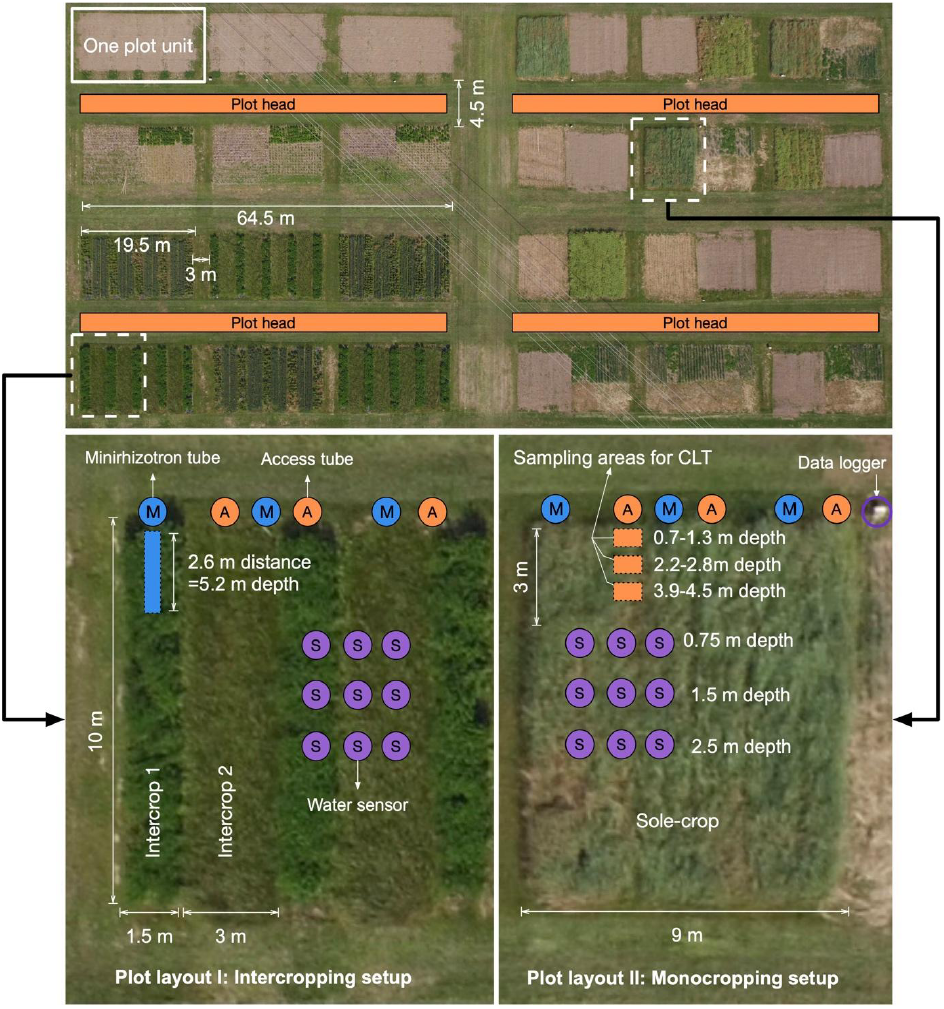
Plot layout at DeepRootLab

**Table 1.**
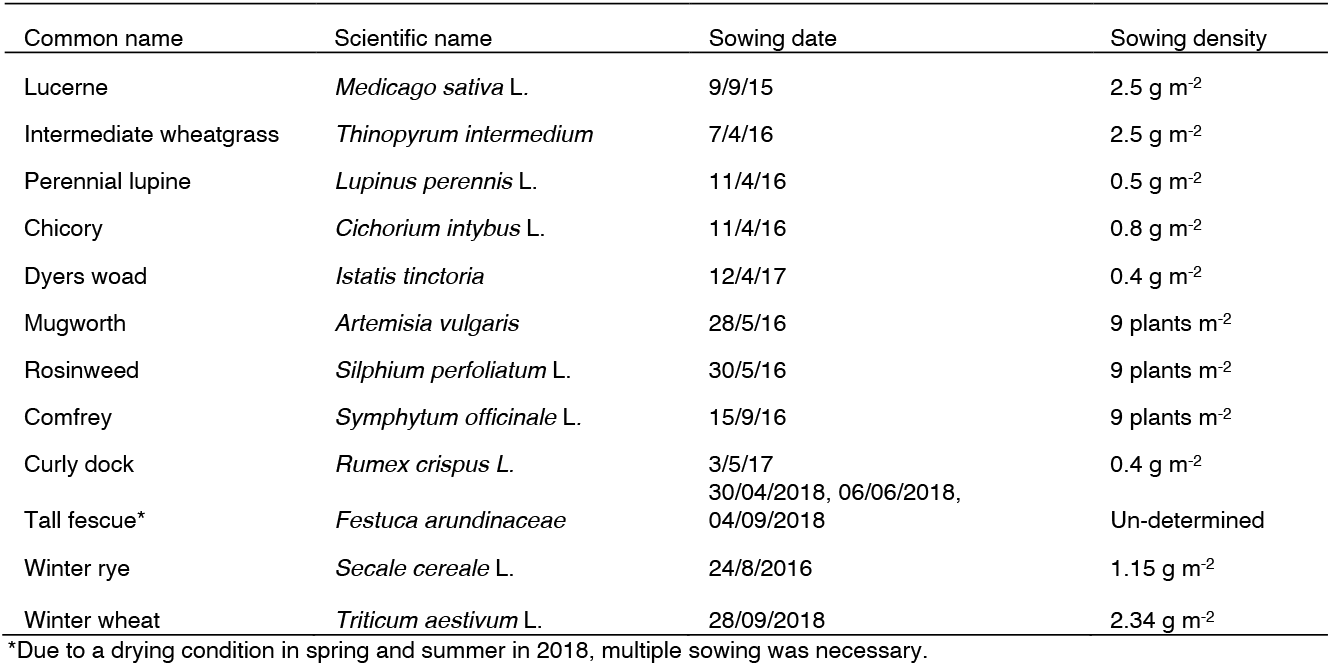
Crop species at DRL.

### Installation of equipment

We used two types of tubes for deep root study (Figure 2), both inserted at an angle of 30º from vertical. The **minirhizotron tubes** are transparent acrylic tubes, 6 m long and have an outer diameter of 70 mm. These tubes were used for non-destructive root measurement with repeated imaging campaigns. The other type is **access-tubes** which are stainless-steel tubes (5.85 m in length, 110 mm in outer diameter). Three cut-out openings were made along the access-tubes, which matched soil depths of 0.7-1.2 m, 2.3-2.7 m and 4.0-4.4 m after installation. **Ingrowth-cores** made of stainless-steel with a volume of 3,931 cm^3^ were made to be filled with soil and inserted into the access-tubes, to allow the study of root growth and uptake processes at the different depths. In total 144 minirhizotron and 144 access-tubes were inserted at a 30º angle vertically at the heads of each plot (**Figure 2**). Inserting the tubes took place from late 2015 to early 2016 via auger drilling.

**Figure 2.**
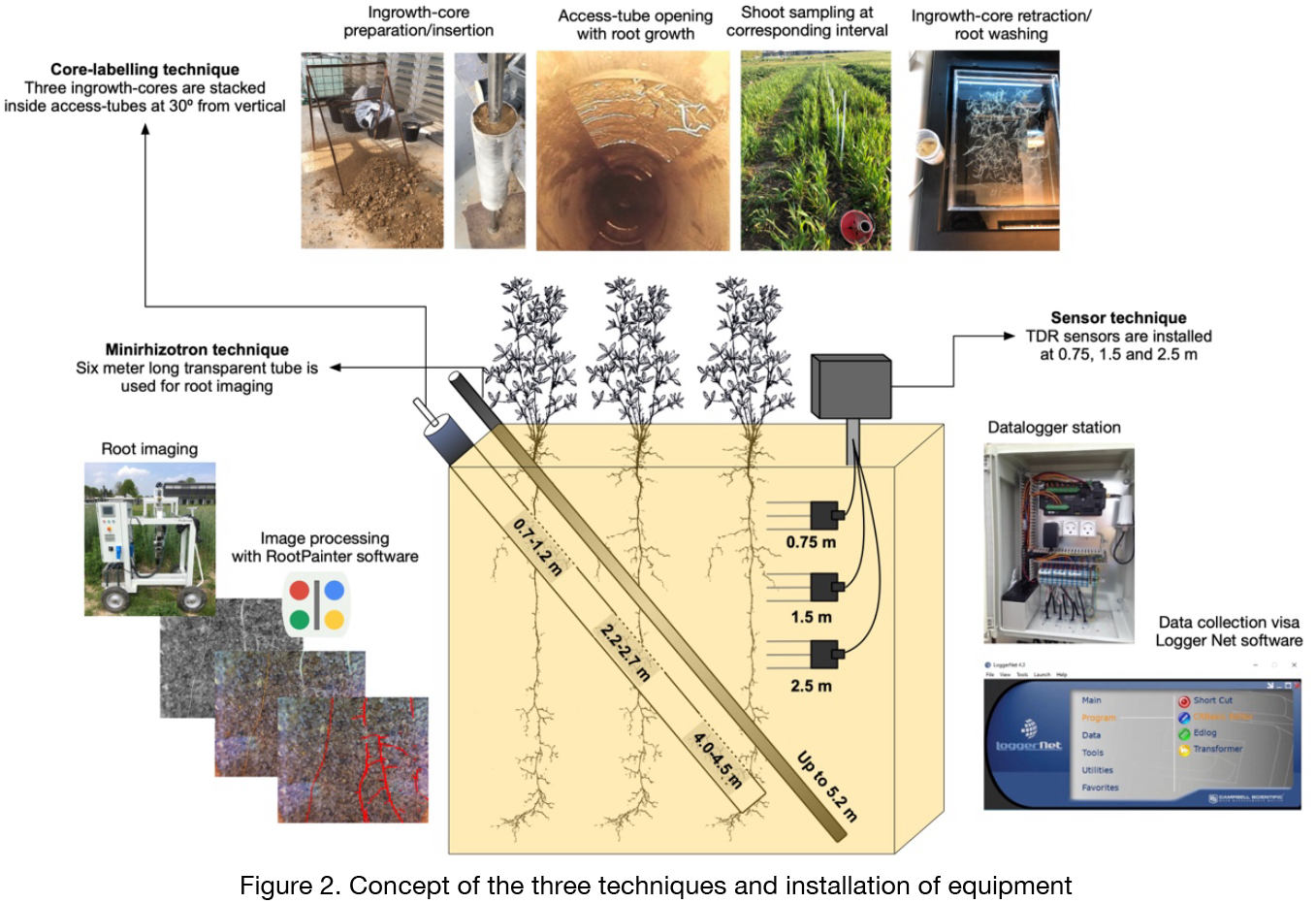
Concept of the three techniques and installation of equipment

Time domain reflectometry (TDR) sensors (Acclima 310S, Acclima, Inc., USA) were installed for monitoring soil volumetric water content (VWC). In selected plots, three replicated water sensors were installed between late 2017 to early 2018 at each of the depths 0.75, 1.5 and 2.5 m, with measurements being recorded on an hourly basis using four CR6 dataloggers (Campbell Scientific, Inc., USA). All TDR sensors were located at least 3 m away from the plot edges (see ‘S’ symbols in **Figure 1)**. The cables were buried at 0.3 m to allow the use of agricultural machinery.

We developed an AI-based software for automated root image analysis called RootPainter (Smith *et al*., 2022). The software features a user-friendly interface that can be easily operated by researchers without experience in Machine Learning. RootPainter utilizes a convolutional neural network (CNN) that is built on the U-Net architecture. After its release, the software was validated for root quantification during destructive sampling (Han *et al*., 2021b).

### Measurement of root growth dynamics

We monitored root growth dynamics of lucerne (2016-2019), intermediate wheatgrass (2016-2019), perennial lupine (2016-2019), mugwort (2016-2019), rosinweed (2017-2019), comfrey (2017-2019), curly dock (2017-2019) and tall fescue (2019-2020) by minirhizotron technique **(Table 2)**. Multispectral images were taken along the minirhizotron tubes at each 0.05 m-length interval by a Videometer lab instrument (Videometer A/S, Hørsholm, Denmark). Multispectral images were obtained by using LED light in five different wavelengths ranging from UVA to violet, amber, red and NIR was adjusted to 365, 405, 590, 660 and 970 nm, respectively. In the results shown, the images were converted to pseudo-RGB images prior to further analysis. The size of each of the images was 40 mm x 50 mm (width x height) consisting of 2048 x 2448 pixels. From the images, **root density** was determined as planar root length density (pRLD) – root length per image area (cm cm^-2^) after training the model using a deep learning software (see results section for detailed description). The **maximum root depth** of each tube was manually determined. For each crop species, we selected one image dataset per year during the crop season. The root depth of each species over accumulative thermal time (ºC days) was computed.

**Table 2.**
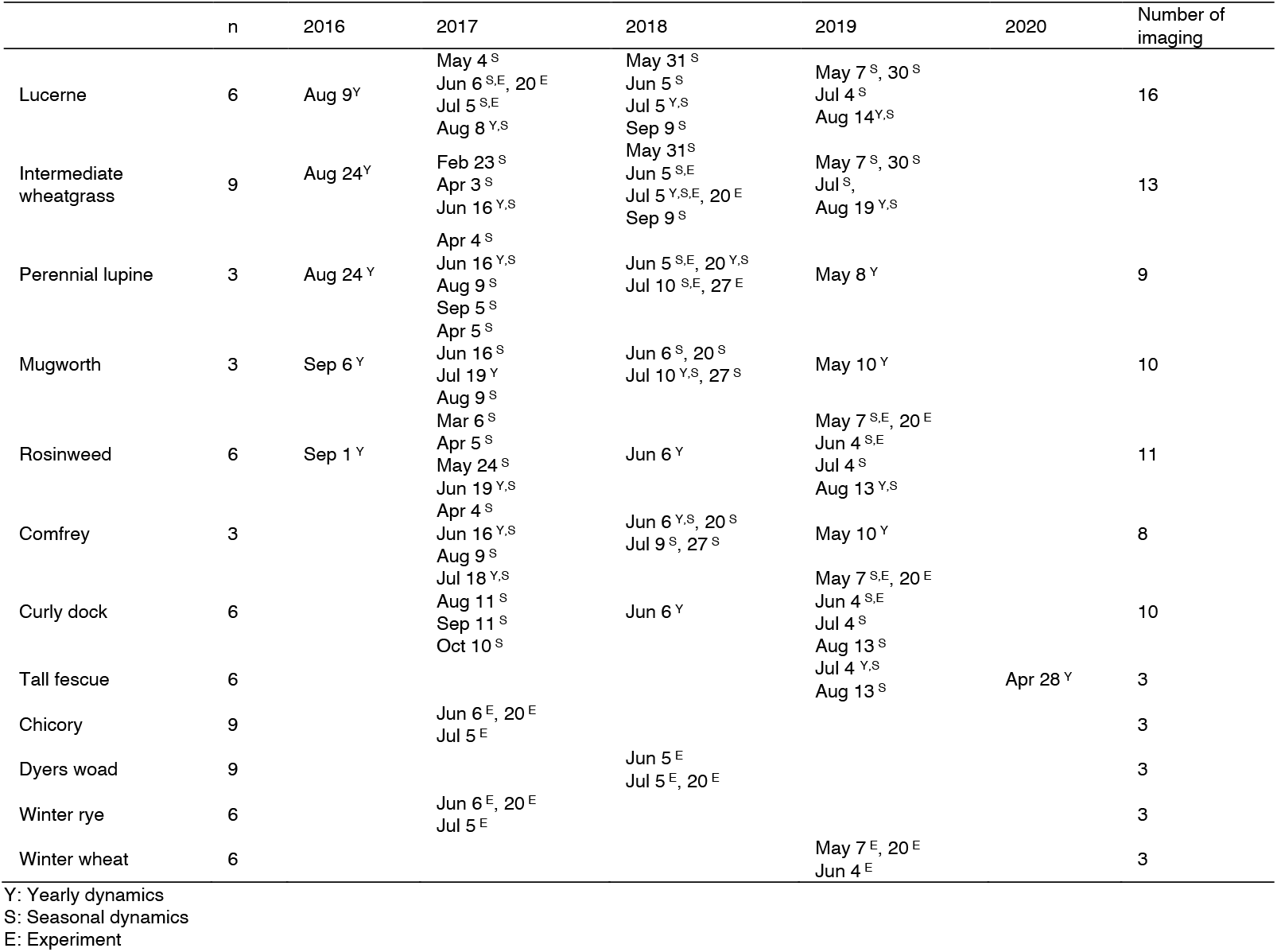
Minirhizotron imaging timeline for 12 crop species at DeepRootLab facility.

**Table 2.** Experimental design for three experiments.

| Experiment |  | Exp 1 | Exp 2 | Exp 3 |
| --- | --- | --- | --- | --- |
| Year/Period |  | 2017<br>May 4 -Jul 4 | 2018<br>May 25-Jul 25 | 2019<br>Apr 1-May 31 |
| Crop species | Early sown perennials | Lucerne (665)* | Mugwort (819) | Rosinweed (1100) |
|  | Late sown perennials | Chicory (450) | Dyers woad (368) | Curly dock (762) |
|  | Annual cereals | Winter rye (315) | Intermediate wheatgrass (768) | Winter wheat (249) |
| Soil depth |  | 0.7-1.2 m | 0.7-1.2 m | 0.0-0.3 m |
|  |  | 2.2-2.7 m | 2.2-2.7 m | 0.7-1.2 m |
|  |  |  |  | 2.2-2.7 m |
|  |  |  |  | 4.0-4.5 m |

\*Days after sowing

### Root activity measurement by core-labelling technique

From 2017 to 2019, we conducted three experiments (Exp 1, 2 and 3) to compare three crop species for deep root growth and nutrient uptake potential **(Table 2)**. Each experiment included at least two perennial species with relatively early sowing dates (lucerne, mugwort and rosinweed) and later sowing dates (chicory, dyers woad and curly dock). In addition, winter cereals or a perennial cereal crop were included for comparisons (winter rye, intermediate wheatgrass, and winter wheat).

We designed an experimental workflow called core-labelling technique (CLT) for measuring deep root uptake (Han *et al*., 2020). Root-free soil medium labelled with tracers was packed inside the ingrowth-cores and inserted into the access-tubes. Ingrowth-cores were installed at two or three depths, matching the cut-out openings of the access tubes. We used various tracers, here we present results obtained with ^15^N as a nutrient tracer. For each ingrowth-core, 275.24 mg of ^15^NH_4_Cl was prepared in solution and mixed with a subsoil medium (see Supp. Table 2 for physical and chemical characteristics) prior to packing the soil into the ingrowth-cores. Tracer concentration determined from the shoot biomass was used as an indicator of root activity.

Ingrowth-cores were inserted at two depth levels in Exp 1 and 2 (0.7-1.2 m and 2.3-2.7 m), whereas four depth levels were investigated in Exp 3 (0-0.3 m, 0.7-1.2 m, 2.3-2.7 m and 4.0-4.4 m), where an extra ingrowth core was installed directly in the topsoil to complement the three deeper placements using the access tubes. Shoot biomass was sampled twice, at the 4^th^ and 8^th^ weeks after insertion of the ingrowth-cores. The dried and powdered samples were analyzed for ^15^N enrichment. For each experiment, root growth of each crop species was monitored using minirhizotrons.

#### Water content measurement by sensors

In this study, soil water content was monitored from June 2018 to June 2020 in three replicated plots for chicory, lucerne and intermediate wheatgrass, and in a single plot for perennial lupin, dyer’s woad and oilseed rape as a pilot trial.

### Statistical analysis

R software (R Development Core Team, 2019) was used for statistical analysis. Linear regression between segmentation and manual counts was performed to compute the R^2^ values as it was affected by increasing annotation time. Root depth dynamics of the eight key species (lucerne, intermediate wheatgrass, perennial lupine, mugwort, rosinweed, comfrey, curly dock and tall fescue) between the years (2016 to 2020) was analyzed using Tukey HSD (P≤0.05). pRLD of the crop species was compared between the growing seasons regardless of the soil depth. More detailed comparisons within seasons in 2017, 2018 and 2019 between the image campaigns were done for four crop species (lucerne, intermediate wheatgrass, rosinweed and tall fescue). For three experiments conducted, pRLD measured by minirhizotrons was compared between the crop species at 0-0.3 m, 0.3-0.7 m, 0.7-1.3 m, 1.3-2.3 m, 2.3-2.8 m and 2.8-4.0 m of soil depth. RLD and **δ**^15^N (‰) were compared between the crop species at soil depths of 0-0.3 m, 0.7-1.3 m, 2.2-2.8 m and 3.9-4.5 m within each experiment. Linear regressions of RLD vs **δ**^15^N (‰) measured from the ingrowth-cores, RLD by ingrowth-cores vs root length by minirhizotron and root length by minirhizotron vs **δ**^15^N (‰) were computed combining the data from the nine crop species used in the three experiments. R^2^ values were computed for each crop species.

## Results

### Data generation capability at DRL

#### Automated root phenotyping

The main data-gathering operation incurred at the facility has been the root imaging via minirhizotron tubes **(Figure 2)**. In our setup, we had an efficiency of acquiring one image per every 5 sec. Consecutive images to 2.3 m to 4.3 m depth per tube were taken. However, depending on season and crop, the imaging depth was often varied. Root growth for most of the crops was monitored using 6 minirhizotron tubes (=6 replicates), except for perennial lupine, mugwort and comfrey which had 3 replicates, and intermediate wheatgrass with 9 replicates. Imaging a tube down to 2.3 m and 4.3 m took 4.5 min and 8.3 min, respectively. Considering the movement and setup of the camera wagon, it was possible to image 36 tubes per day. This allowed multiple imaging campaigns per treatment per season throughout the facility.

The raw image format (Hips) from the camera was converted to a more common JPEG format. In our estimate, 5.2 sec per image was required for this process. We converted 61,471 images which involved approximately 60 hours. Afterwards, we spent approximately 10 hours training the dataset using the RootPainter software, which had an efficiency of 0.6 sec per image for inference.

Therefore, in our estimate, 11.8 sec per image was spent on image capture, processing, and model training for segmentation. In total, we processed 61,471 images acquired over 5 years (2016 to 2020). This required approximately 200 hours for the entire process which is approximately 25 days-which is equivalent to 5 working days per year.

#### Root activity determination at DRL

In total, 36, 63 and 96 ingrowth cores were installed in Exp 1, 2, and 3, respectively. In our estimate, three workdays were needed to complete the labelling and packing of 36 ingrowth-cores. Insertion/retraction of ingrowth-cores into/from the access-tubes required ideally three persons and 1-2 days of operation time depending on the weather conditions and occurrence of mechanical disturbance, e.g. ingrowth-cores jammed inside the access-tubes.

#### TDR sensor recording

TDR sensors’ data were recorded hourly and data withdrawal was done manually using the Logger Net software. Downloaded “dat” files were used to extract VWC data. Soil water content data were averaged daily for this study.

### Automated root segmentation

#### Model performance

We report the performance of a root segmentation model trained on the root images using the RootPainter software. Manual counts on 200 random images were correlated against the automatic segmentation over the period of annotation/training **(Figure 3)**. After 200 minutes of annotation, R^2^ reached 0.81. After around 10 hours of corrective annotation, we found an R^2^ of 0.91 with the manual root counted data, showing high root detection accuracy.

**Figure 3.**
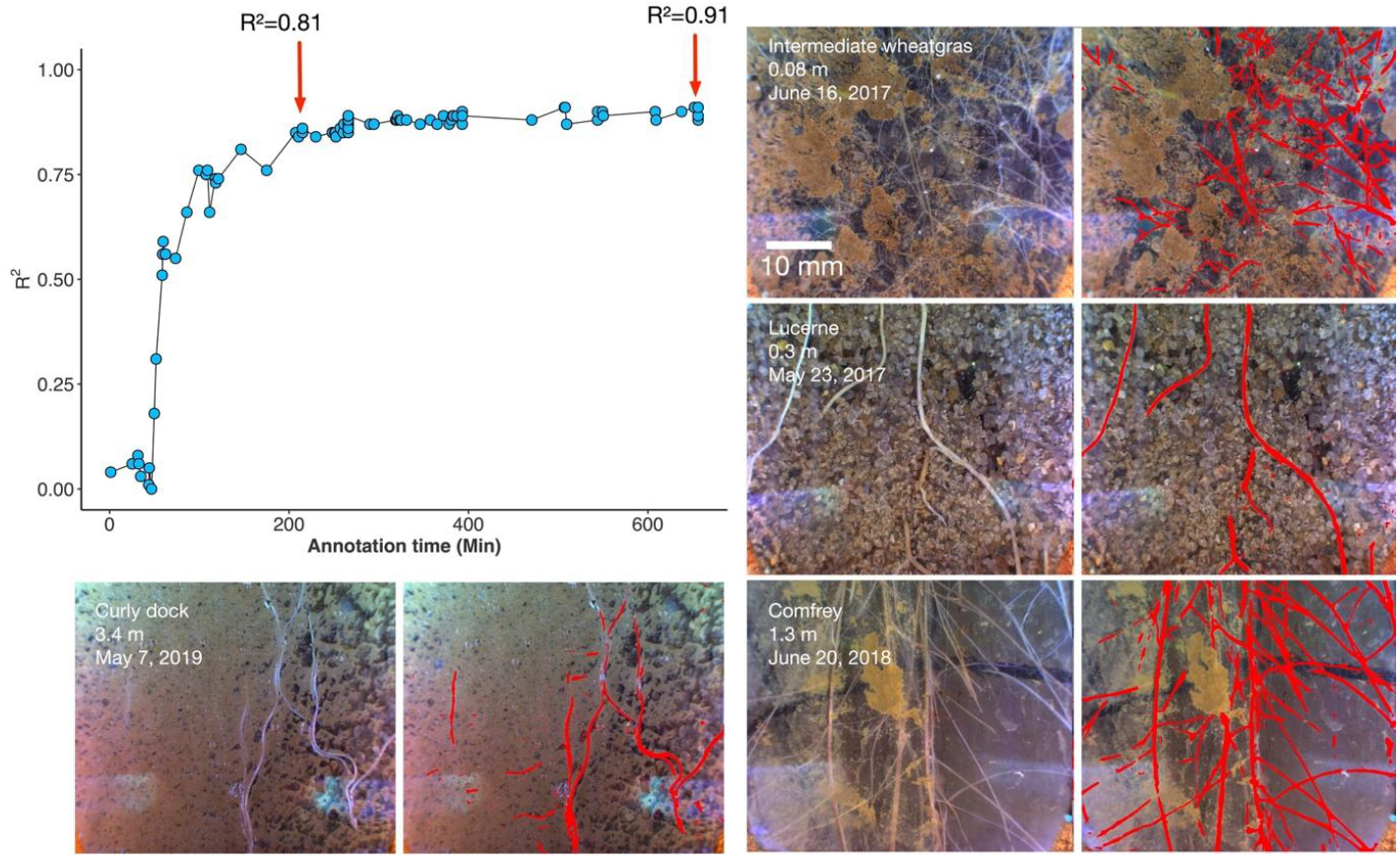
R-values derived from linear regression between segmentation and manual counts as a function of annotation time. Comparative view on original (left) and segmented (right) images.

#### Training process

We modified the established training protocol by Smith *et al*. (2022). The original protocol advises annotation on 6 examples before initiation of corrective annotation but we did not see a clear improvement in prediction after 6 images **(Figure 4A**,**B)**. Therefore, we initiated corrective annotation after 16 clear examples when we observed roots being segmented with a clear shape **(Figure 4C,D)**. False positives on non-root objects such as the scratches on minirhizotron tubes **(Figure 4E)** and water bubbles **(Figure 4F)** prevailed throughout the training process. We focused on making corrections to the false positives. As a result, the model started to exclude the artefacts accurately after the 467th loaded image.

**Figure 4.**
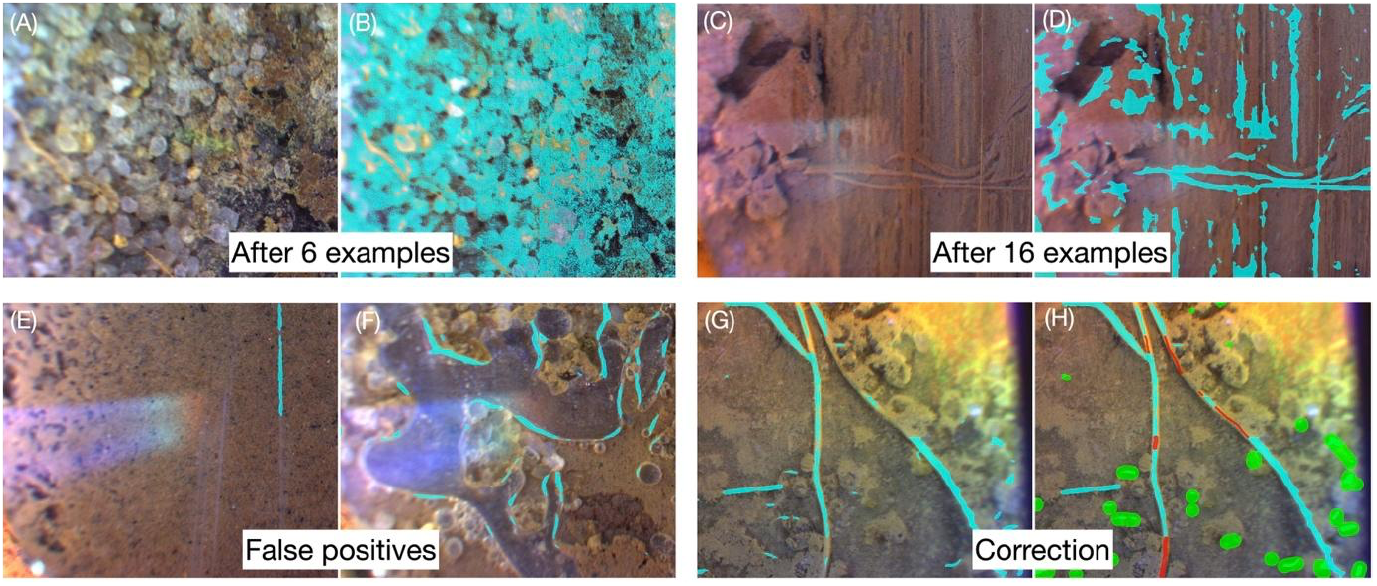
Prediction after six (A, B) and 16 annotations (D, E); false positives – scratches and water bubbles (E,F); corrective annotation to connect the gaps in root segmentation and remove false positives. Red brush, green brush were used for foreground and background. Prediction is shown in aqua blue.

We also occasionally observed false negatives i.e., roots not predicted as roots after corrective annotation was initiated - for example, one single root was segmented as separate parts (Figure 5G: at 162^nd^ image). Whenever applicable, we connected the gaps between the separate segmentations as one single root (Figure 4H). We suspect these efforts improved the predictions substantially towards the end of training.

**Figure 5.**
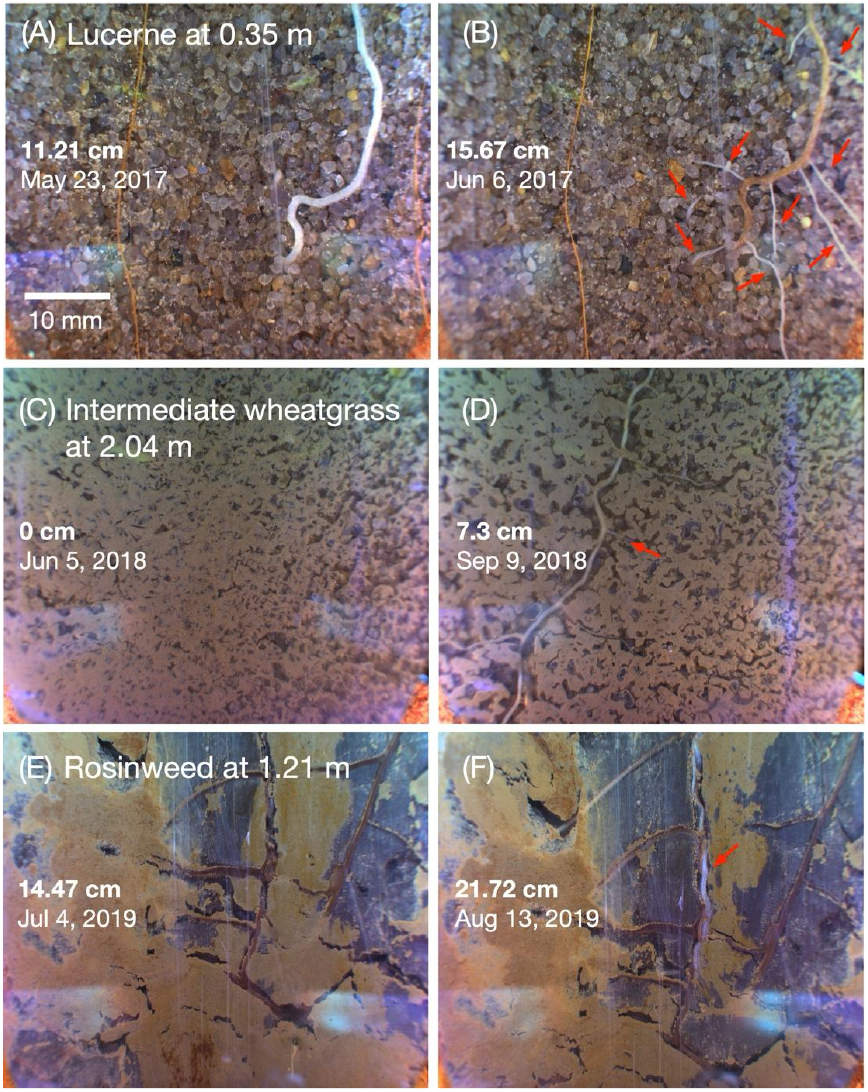
Visually captured seasonal root growth dynamics. Lateral branching of lucerne roots at 0.35 m taken on May 23 (A) and Jun 6 (B) in 2017. Penetration and lateral branching of intermediate wheatgrass roots into new root depth at 2.04 m taken on June 5 (C) and Sep 9 (D) in 2018. Penetration of rosinweed roots into pre-existing root depth at 1.21 m taken on July 6 (E) and Aug 13 (F) in 2019. Arrows indicate new root growth. Root length (cm) was derived from segmentation.

Most importantly, the trained model accurately predicted root growth when (1) old branches produced laterals **(Figure 5A,B)**; (2) roots penetrated into a new depth **(Figure 5C,D)**; (3) roots penetrated into a depth with old roots **(Figure 5E,F)** that provided quantitative information on the increment of root density (in cm) between two imaging intervals.

### Root depth and root penetration rate

We determined root depth dynamics of the eight perennial crop species **(Figure 6)**. Overall, there was a clear trend of increasing average root depth over the years by all measured crop species. The root penetration rate computed by regression analysis between root depth over the thermal time (ºC days) was slowest for lucerne (0.07 mm ºC days^-1^) followed by perennial lupine (0.08 mm ºC days^-1^), intermediate wheatgrass (0.1 mm ºC days^-1^), mugwort (0.2 mm ºC days^-1^), tall fescue (0.24 mm ºC days^-1^), curly dock (0.26 mm ºC days^-1^), rosinweed (0.29 mm ºC days^-1^) and comfrey (0.38 mm ºC days^-1^). Maximum root depth - the deepest point of root observation was shown by curly dock and rosinweed in 2019 at 3.8 m, followed by tall fescue (3.5 m), intermediate wheatgrass and comfrey (2.9 m), lucerne (2.7 m) and mugwort (2.6 m).

**Figure 6.**
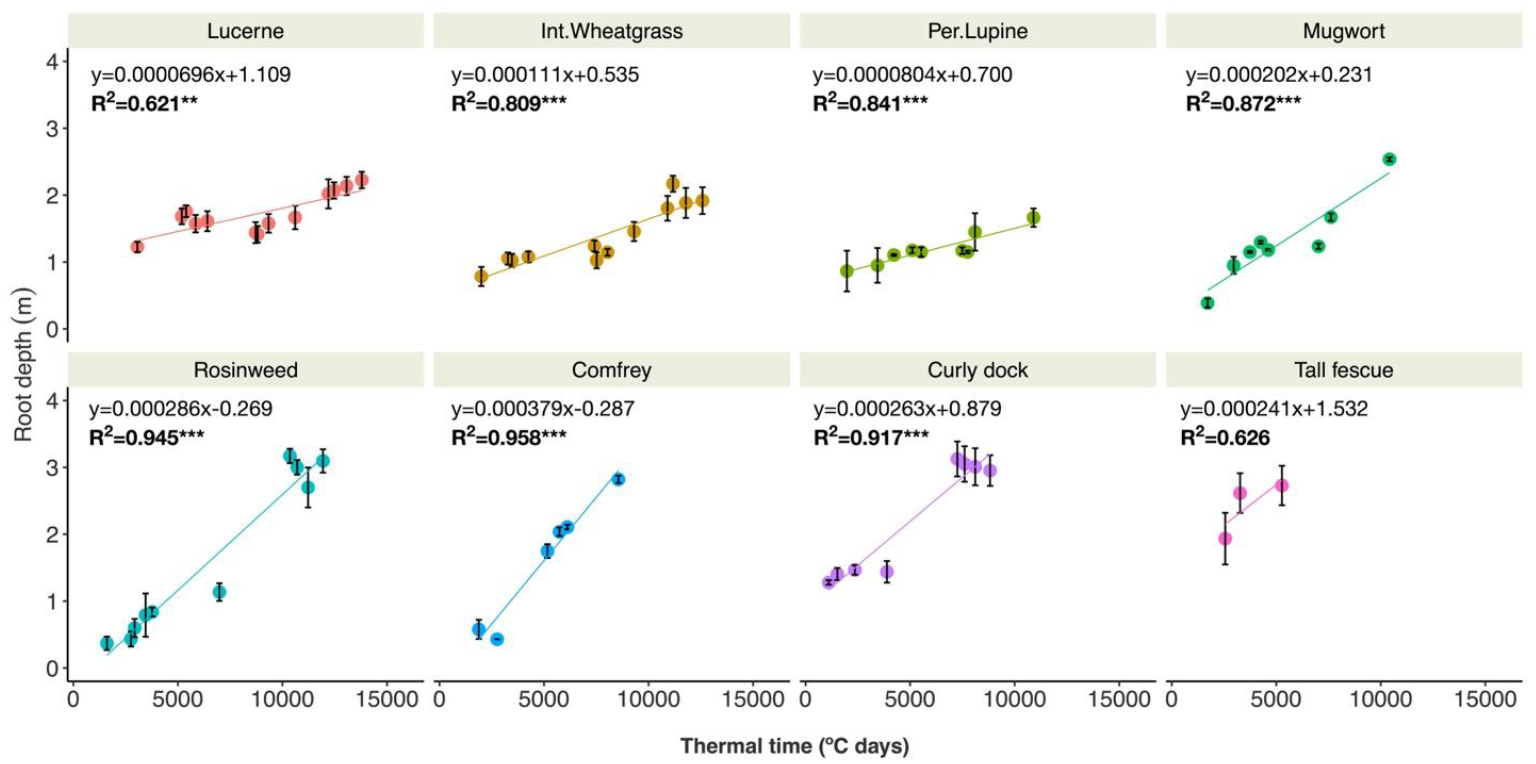
Root depth dynamics of perennial species at DeepRootLab facility measured over thermal time (ºC days) from 2016 to 2020. Mean and SE (± one) is shown plotted over linear regression lines.

### Yearly dynamics in root density

We compared the root density (measured as planar root length density, pRLD (cm cm^−2^) of 8 perennial species between crop seasons at each 0.5 m interval to 4.0 m of soil depth (**Figure 7)**. Overall, root length generally increased as the crops grew older. The effect was most pronounced below 0.5-1.0 m of soil depth, except for mugwort and tall fescue. Intermediate wheatgrass exhibited greater root length in its 2nd growing season (in 2017) especially in the upper soil layer (0-0.5 m) but its root density decreased over the following two years. Perennial lupine did not show any effect of crop age on root length. All crop species revealed deeper rooting from the first to the final point of observation.

**Figure 7.**
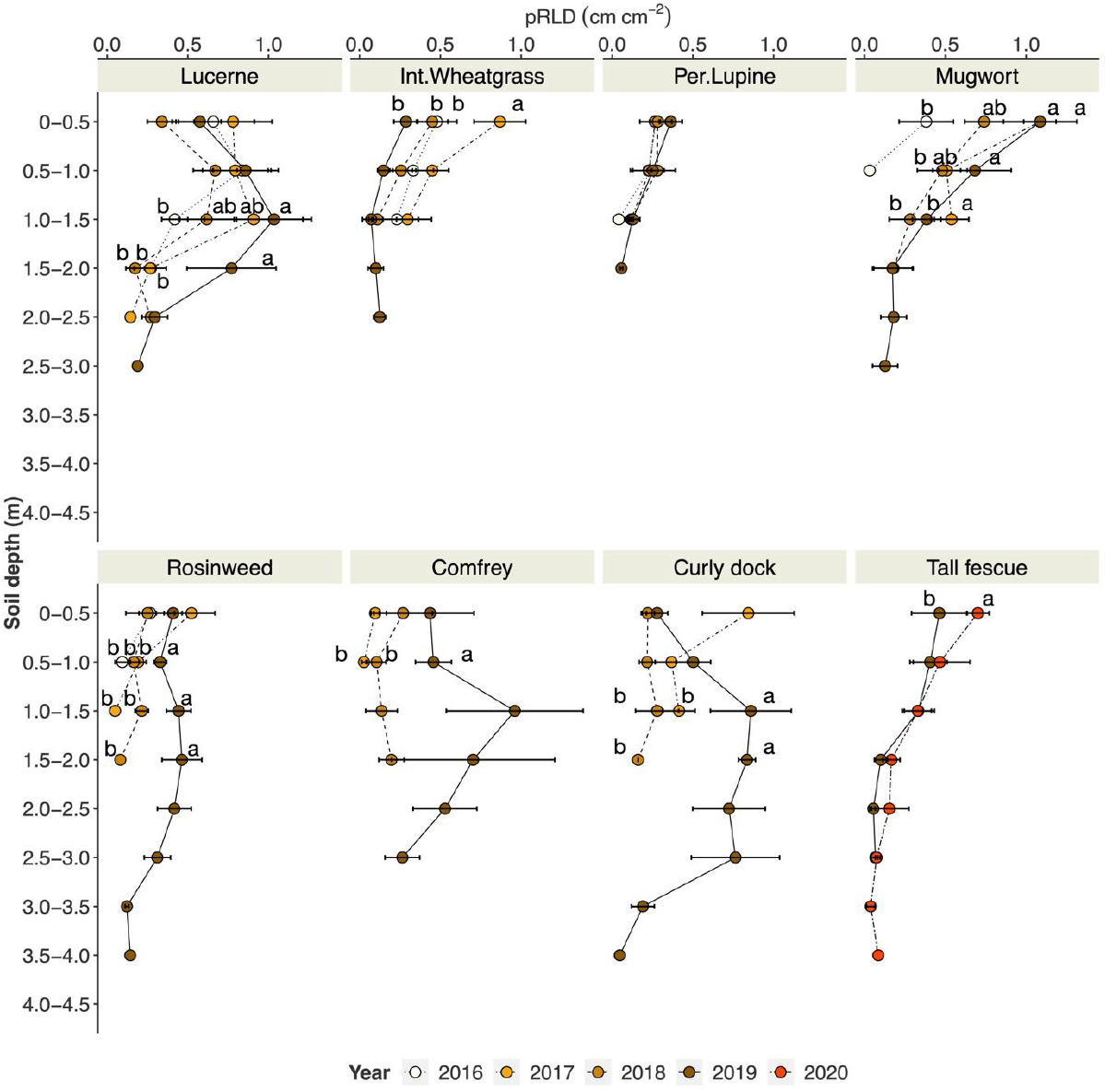
Yearly root growth dynamics of lucerne, intermediate wheatgrass, perennial lupine, mugwort, rosinweed, comfrey, curly dock, tall fescue expressed as planar root length density (pRLD; cm cm^-2^). Small letters indicate significant differences between the year (Tukey HSD; P≤0.05).

### Seasonal root growth dynamics

We compared the root density of lucerne, intermediate wheatgrass and rosinweed at each depth-level between the two observation times within the crop seasons in 2017, 2018 and 2019 **(Figure 8)**. Root density of lucerne remained unchanged in 2017 between the observation points, but in the following two years, an increase in root density was shown below 1.0-1.5 m soil depth. In 2019, a significant decrease in root density was observed in the topsoil. Intermediate wheatgrass showed little effect of measurement time, but with a tendency to declining root density later in the season. Rosinweed exhibited an increase in root density within the season in 2017 (0-0.5 m) and 2019 (0.5-1.0, 1.0-1.5, 1.5-2.0 m).

**Figure 8.**
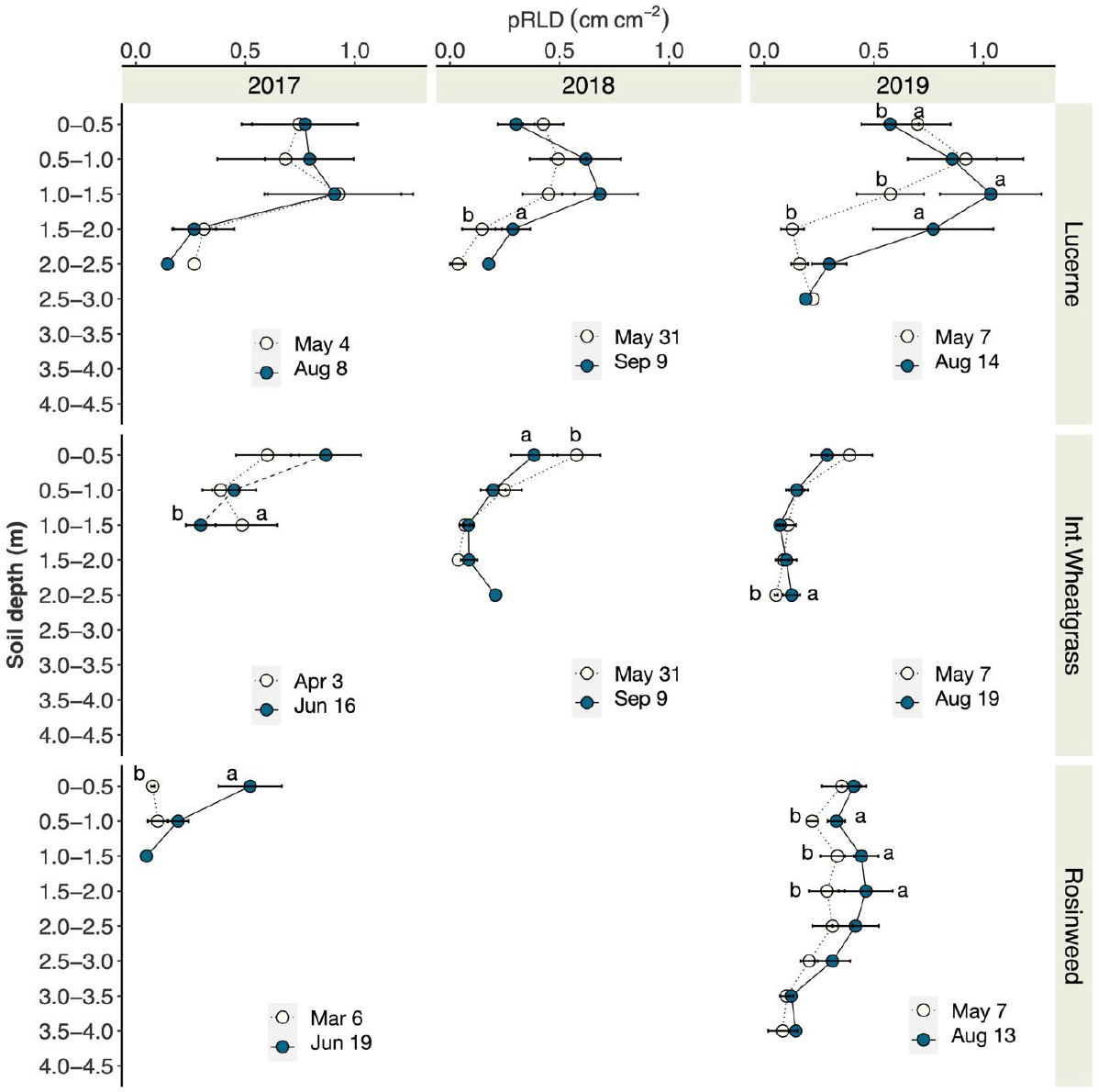
Seasonal root growth of lucerne, intermediate wheatgrass, rosinweed and tall fescue measured in 2017, 2018 and 2019 expressed as planar root length density (pRLD; cm cm-). Small letters indicate significant differences between the image campaign (Tukey HSD; P≤0.05).

### Crop comparison for deep root development

Using the minirhizotron technique, we observed a clear difference in root depth among the compared crop species **(Figure 9)**. In Exp 1, chicory showed the deepest roots at 2.2-2.3 m depth followed by lucerne (2.1-2.2 m) and winter rye (1.4-1.5 m). In general winter rye had the lowest and chicory the highest root density along the soil depth profile. Root density of chicory was greater than lucerne in the upper soil layers (0-0.7 m).

**Figure 9.**
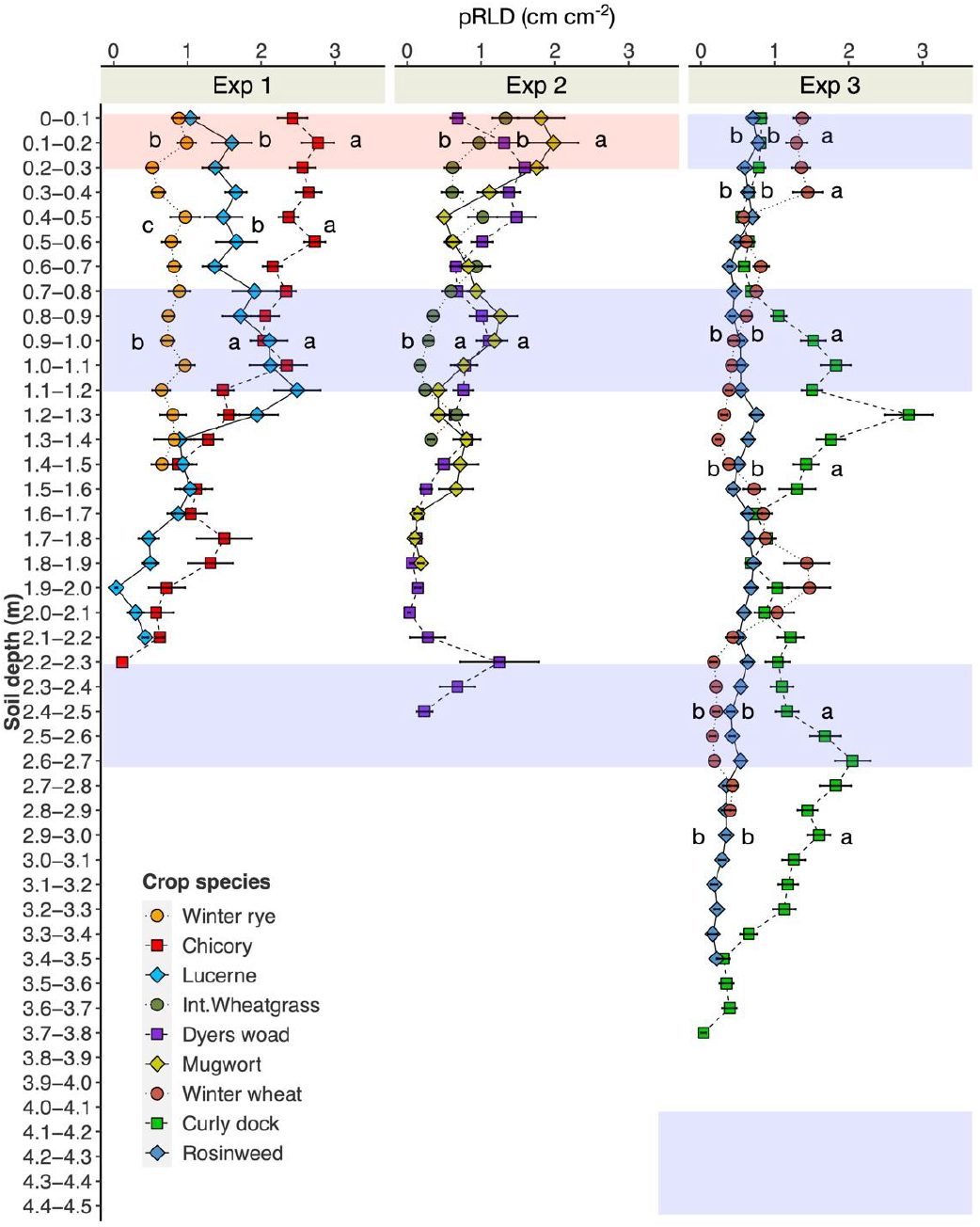
pRLD (cm cm^-2^) measured at Exp 1, 2 and 3 by minirhizotron imaging and segmentation. The images were acquired 8 weeks after Ingrowth-core insertion. Purple strips indicate the depth range for ingrowth-core measurement. No ingrowth-core was inserted in the white and red strips. Small letters indicate significant difference between the crop species within the depth range (red, white and purple) at each experiment (Tukey HSD: P≤0.05).

In Exp 2, dyers woad showed the deepest roots (2.4-2.5 m), followed by mugwort (1.8-1.9 m), and intermediate wheatgrass (1.3-1.4 m). In the topsoil, root density of mugwort was greater than other crops. There was some variability with depth, but in general, intermediate wheatgrass showed lower root densities than dyers woad and mugwort.

We found roots of curly dock to 3.8 m depth in Exp 3. Winter wheat and rosinweed grew roots to 2.8-2.9 m and 3.3-3.4 m, respectively. Except in the topsoil layer, curly dock exhibited a higher root density compared with rosinweed and winter wheat.

### Crop comparison for deep root activity

We compared deep root activity of three crop species by inserting ingrowth-cores packed with ^15^N-labelled soil in Exp 1, 2 and 3. Two parameters were measured in all experiments, namely, RLD (cm cm^-3^) from the root extraction from the soil in the ingrowth-cores **(Figure 10A)** and **δ**^15^N (‰) concentration measured from aboveground biomass **(Figure 10B)**.

**Figure 10.**
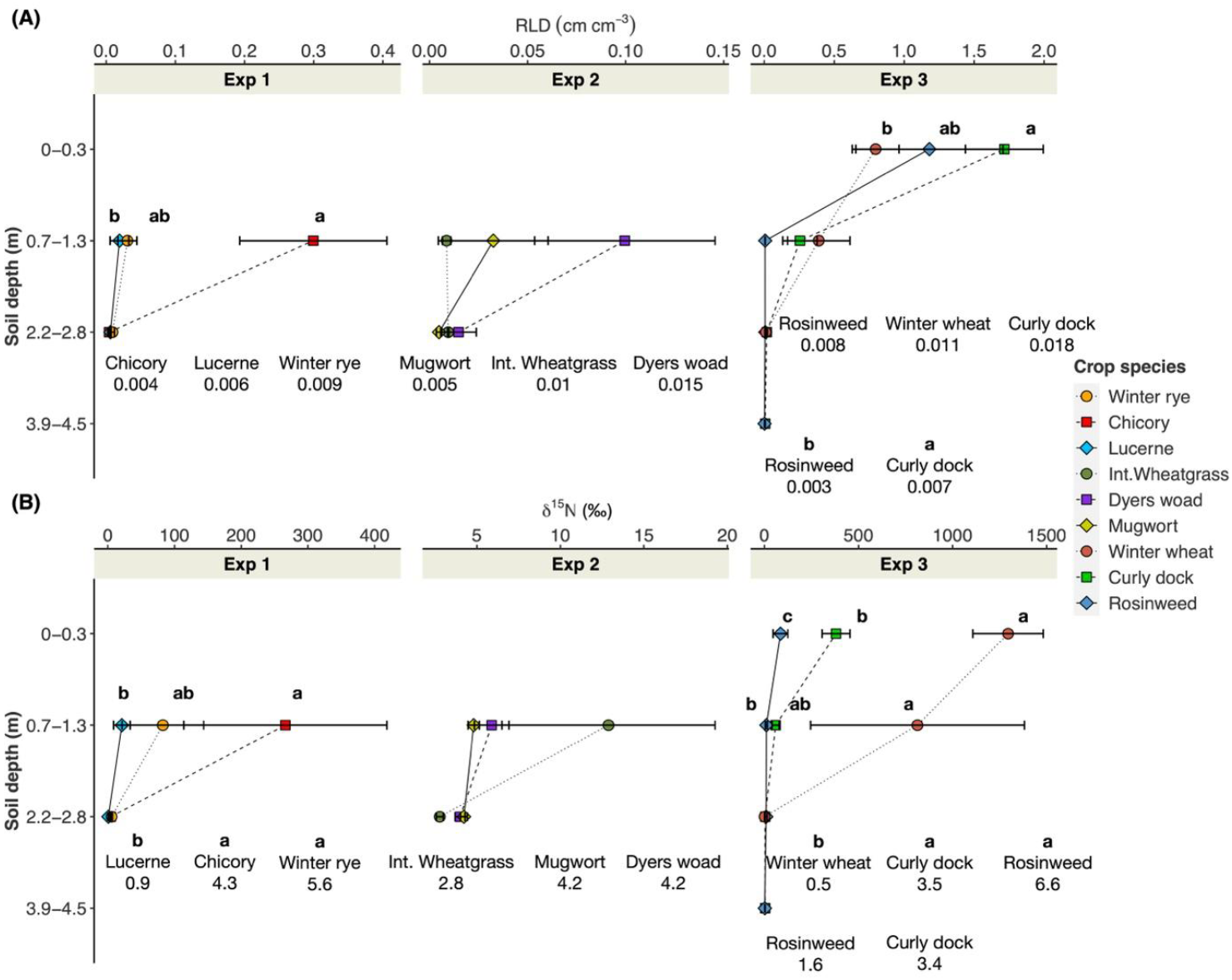
Root length density (A; RLD; cm cm^-3^) and **δ**^15^N (per mill) concentration measured in Exp 1, 2 and 3. Small letters indicate significant differences between the crop species within the soil depth (Tukey HSD: P≤0.05). For 2.2-2.8 m and 3.9-4.5 m depths, RLD and, RLD and **δ**^15^N values are shown in numbers.

In Exp 1, the ingrowth-cores were inserted at 0.7-1.2 m and 2.2-2.7 m depth-levels. At 0.7-1.2 m, chicory exhibited greater RLD and **δ**^15^N compared with lucerne, while intermediate results were shown by winter rye. No difference in RLD was found between the crop species at 2.2-2.7 m. However, **δ**^15^N of winter rye and chicory was greater than lucerne at this depth.

Ingrowth-cores were inserted at the same depth-levels in Exp 2. No meaningful difference in RLD and tracer concentration was found in Exp 2. Overall values of RLD and **δ**^15^N were low, especially at 0.7-1.2 m.

In Exp 3, the following four depth-levels were investigated: 0-0.3 m, 0.7-1.2 m, 2.2-2.7 m and 4.0-4.5 m. In the topsoil layer, the differences between the crop species in RLD and **δ**^15^N were not consistently shown. RLD of winter wheat was low, and significantly lower than for curly dock, but **δ**^15^N of winter wheat was substantially greater than for both other crops in topsoil layer. No difference in RLD was found at 0.7-1.2 m, yet winter wheat exhibited greater **δ**^15^N compared with rosinweed. At 2.2-2.7 m, greater **δ**^15^N was found for rosinweed and curly dock compared with winter wheat, whereas no difference in RLD was found. RLD of curly dock was greater than rosinweed at 2.2-2.7. A similar trend was found for **δ**^15^N at 4.0-4.5 m, however, the difference was not significant.

### Validating measurements

Regression analysis between the root density found inside the ingrowth-cores and tracer uptake (δ^15^N) measured from aboveground biomass showed a significant relationship (P≤0.001) with R^2^ values of 0.67 when all data from the three experiments were combined (Figure 11A). When the same analysis was performed for each crop species, it revealed strong correlations for chicory (R^2^=0.97; P≤0.001) and winter wheat (R^2^=0.92; P≤0.001). Except for lucerne, dyers woad, mugwort, the relationship between RLD and **δ**^15^N was found to be significant for the rest of crops.

**Figure 11.**
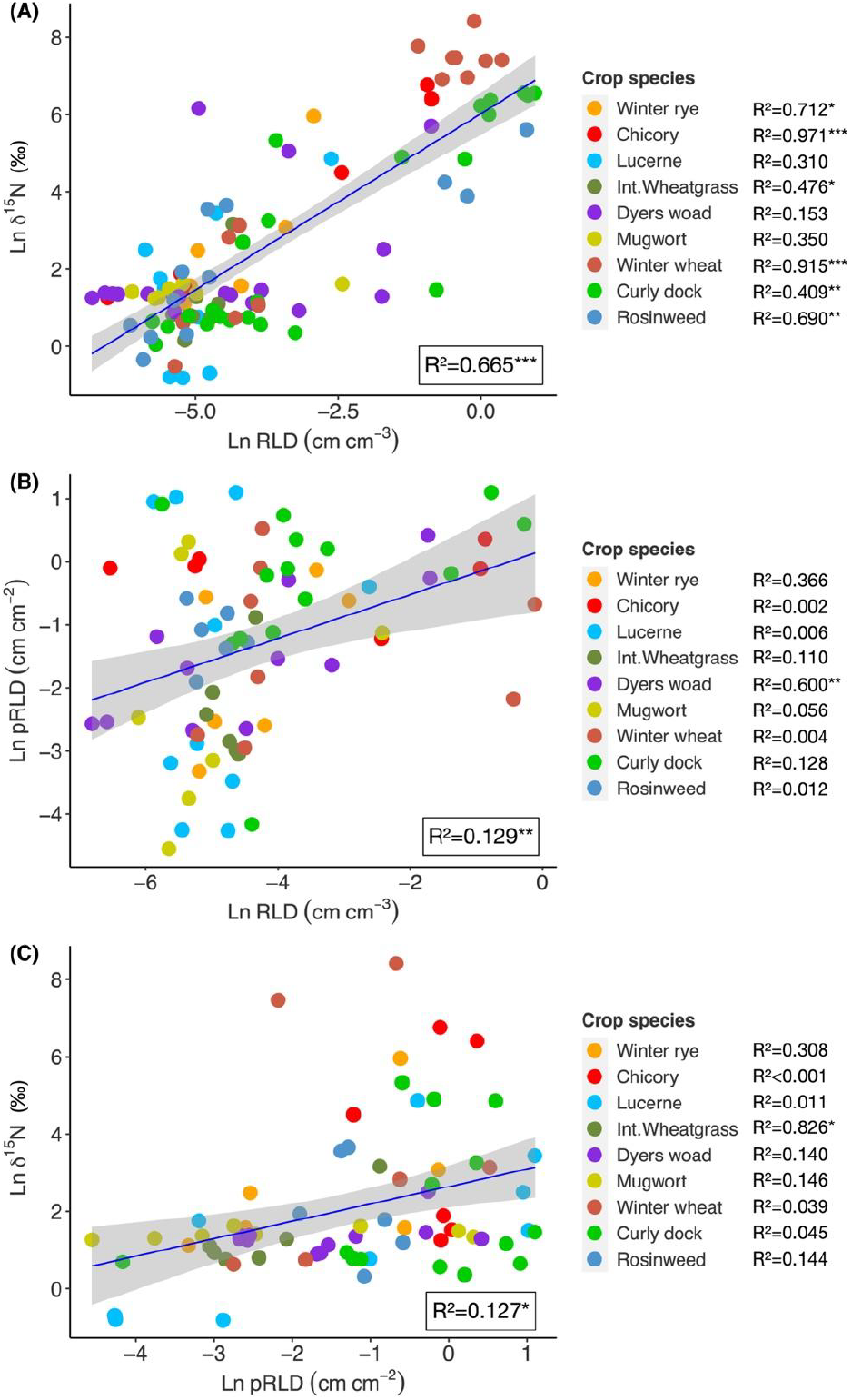
Linear regressions between root-length density (RLD) inside ingrowth-cores and **δ**^15^N concentration (A: RLD vs.δ^15^N); RLD and pRLD measured by minirhizotron (B: RLD vs. pRLD); root length measured by minirhizotron and **δ**^15^N concentration (C: pRLD vs. **δ**^15^N). R^2^ values as a combined dataset for each experiment were shown inside the plots, whereas R^2^ values for individual species are shown in legends.

We performed linear regression analysis between root density found inside the ingrowth-cores and root density measured by minirhizotron imaging (Figure 11B).

The relationship was found to be significant (P≤0.01), however, with a small R^2^ values (0.13). When performed on each crop species, only dyers woad showed a significant relationship with a moderate R^2^ value (0.6; P≤0.01).

Similarly, a weak but significant relationship (R^2^=0.13; P≤0.05) was found between root density observed by minirhizotron and tracer uptake (δ^15^N; Figure 12C). Only intermediate wheatgrass exhibited a significant relationship between these two measurements (R^2^=0.83; P≤0.05*).

**Figure 12.**
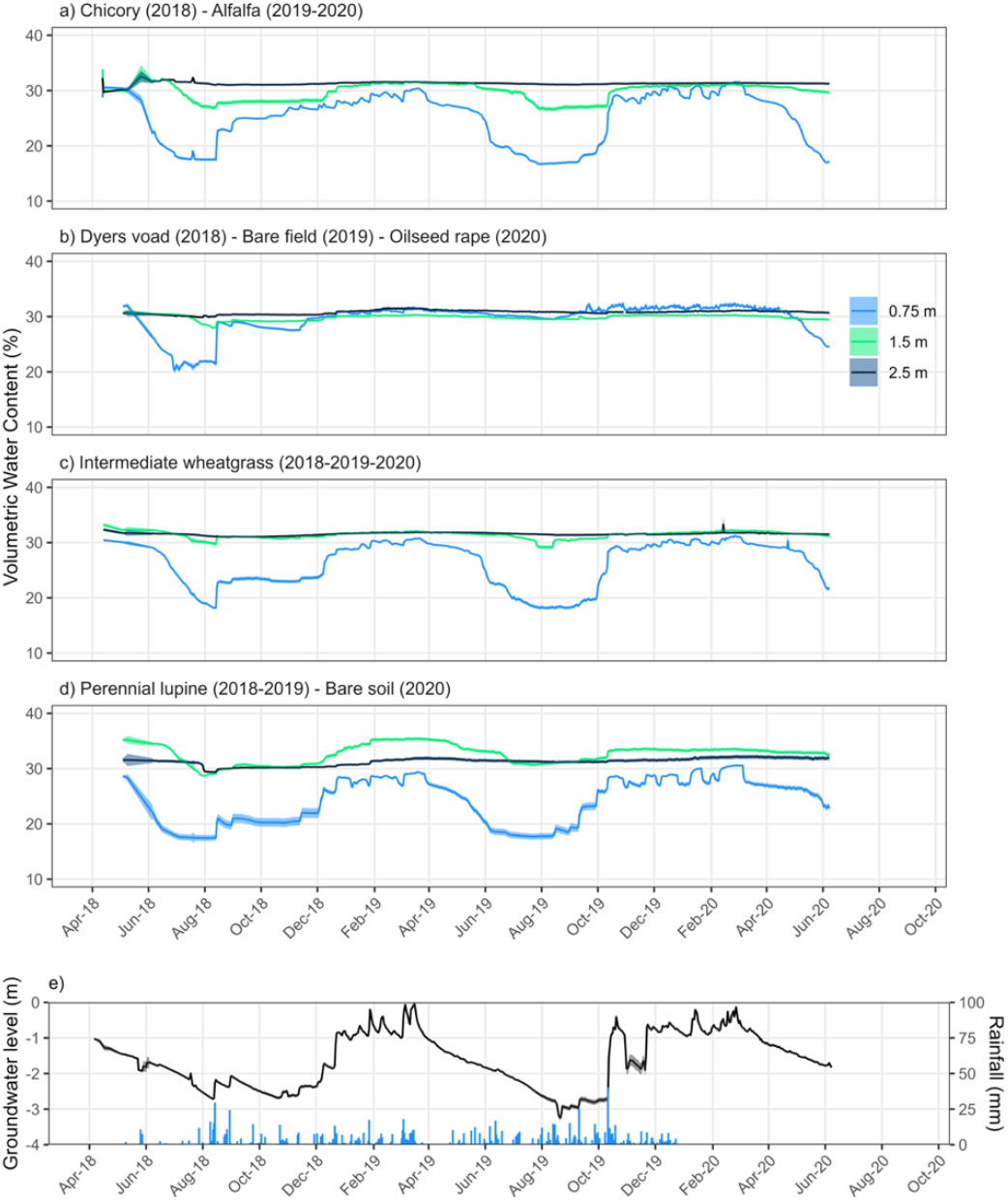
Soil volumetric water content evolution at 0.75 m (light blue), 1.5 m (green) and 2.5 m (dark blue) soil depth under four different crop rotations (a-b-c-d). Crop rotations are given in the figure’s titles. Daily average value with standard error (shaded area). Number of sensors per soil depth are: n_a_= 9, n_b_= 3, n_c_= 9 and n_d_=3. Daily groundwater level evolution (dark line) and rainfall pattern (bars) over the three seasons of cultivation (e). Groundwater level data were averaged over 4 wells and standard error are presented with shaded area. Data covers three season of cultivation from 04-2018 until 10-2020. Note: Lucerne and intermediate wheatgrass data 2018-2019 were already published in Clement et al (2022).

### Water measurement

TDR sensors gave us accurate measurements of the soil water content evolution over time. Soil water content followed a wetting and drying cycle, associated with seasonal differences in crop growth and rainfall patterns (Figure 13). These differences also influenced the groundwater level which is characterized by relatively high “standing” groundwater level in winter and low groundwater level (i.e. > 2.5 m) during the summer (Figure 13E). Over the three growing seasons, all crops were able to deplete water from the 0.75 m soil layer, with values reaching close to wilting point (i.e. 17% VWC) (Figure 13A-D). However, the rate of water depletion at this depth differs among crops. Perennial lupine, chicory and lucerne quickly depleted this soil layer, which almost all its available water used by early June. In contrast, slower rates of water depletion were observed for intermediate wheatgrass and dyers woad with soil water content reaching a value close to wilting point in July (Figure 13B-C). Along with these results, perennial lupine, chicory and lucerne depleted substantially more water from the 1.5 m soil depth compared to intermediate wheatgrass and dyers woad. Indeed, at this soil depth, perennial lupine, chicory and lucerne induced approximately a 5-6% drop in soil water content whereas intermediate wheatgrass and dyers woad induced smaller changes of approximately 2-3%-VWC. However, no crops depleted this soil layer down to a value close to the wilting point. At 2.5 m soil depth, only perennial lupine induced a noticeable decrease of soil water content of 2 % VWC whereas no water use from this layer was shown for any of the other crops.

## Discussion

We went through three major steps to establish the facility; (1) installation of new equipment (minirhizotron and access-tubes, water sensors, data loggers); (2) validating experimental procedures for root activity detection (3) streamlining the root quantification procedure using minirhizotron technique coupled with deep learning-based segmentation.

### Challenges for installation in deeper soil layers

For both types of tubes, i.e. minirhizotron and access-tubes, we faced challenges for insertion due to relatively high clay content (<20 %; see Table 2) at the study site as well as high water table (approx. 1 m) at the time of installation during the winter and early Spring in 2016-2017. The long tubes also - -strongly increase the friction while installing them into the holes after drilling. For minirhizotron tubes, therefore, it was unavoidable to make the holes larger than the tube diameter. This caused a gap between the upper soil surface and the tubes. We attempted to offset this issue by attaching raise-blocks at the bottom side of each tube to ensure better soil-tube contact at the upper side of the sloping tubes. The remaining gaps were filled with sand materials. However, this caused an artificial background on many images with the added sand, and for some tubes, the gaps were not filled (see Figure 5 and Figure 6). This might have caused a low-resistance path leading to an artificial increase in root growth (Rewald & Ephrath, 2013; and references therein). However, the minirhizotron tubes were inserted under the same protocol across the platform for a short period of time. Therefore, it is unlikely that the differences in deep root density between the crop species and observation timing were artefacts. For the metallic access-tubes, we tackled this insertion issue by using sonic-drilling – in which the resonating energy from a high-frequency oscillation (e.g. 150 Hz) in the drill head ensures better penetration into the deep soil layers. After installation, we observed occasionally that some soil from upper opening areas fell into the tubes -which may have created an artefact for root growth by reducing root-soil contact.

### High-throughput root measurement using root image analysis

One of the main advantages of the minirhizotron method is its capacity to measure gross root growth without interfering with the previously measured--area/depth. We have further automated image capture by installing a programmable logic controller (PLC) which generates file names with useful meta-data such as the plot number, location of camera (in terms of cable length projected) and timing of image capture as illustrated by Svane *et al*. (2019a). This enabled us to collect a large amount of data and process the images without much manual effort (e.g. re-naming of the files).

Integrating the automated image capture with deep learning-based analysis, we achieved a high efficiency of root data acquisition (approximately 12 seconds per image). This certainly is a substantial improvement compared to manual root counting--and annotation in terms of time-saving. For example, around 1 to 1.5 hours of annotation time was required for 100 cm^2^ of root image area (Ingram & Leers, 2001), which now can be done in 1 minute using our approach (even including the data capture and image processing time).

Our results showing the relationship between annotation time and root length correlation inform the RootPainter community on the trade-offs of extended annotation for minirhizotron segmentation models. We found a large portion of model improvements to happen within the first 200 minutes, with extended annotation continuing to-provide improvements but with diminishing returns (see Figure 4). These continuing improvements indicate that extending RootPainter annotation and training time past the initial period of rapid improvement can help improve segmentation accuracy and overcome artifacts in challenging heterogeneous field root datasets. Some of our images captured the new root growth between the measurements taken at relatively short intervals. For example, lateral branching of lucerne roots at 0.35 m within two weeks was observed (Figure 6A,B), which was successfully captured by the trained model showing the increase in root length from 11.21 cm to 15.57 cm per image. It was intriguing to see the rapid colour change of the main root from white to brown upon proliferation. Also, the imaging captured root growth of intermediate wheatgrass into depth, growing below 2 m within the growing season. This new occupancy of roots was observed between the measurements taken in Jul and Sep in 2018. Here, we show the images from Jun and Sep, 2018 which better represent the location of imaging (Figure 6C,D). This was also accurately determined by the trained model with the root length increasing from 0 cm to 7.3 cm. We were able to capture newly grown roots grown into the area at 1.21 m where dark roots of rosinweed were present from the growth in previous seasons (Figure 6E,F). As a result, our approach was sufficiently accurate to capture the within-season variation (May to Aug in 2017, 2018 and 2019) in root density of crop plants (see Figure 9), which makes consistent observation possible leading to high-time resolution dataset.

### Tracer-based root activity measurement

Pre-installation of access-tubes, insertion of ingrowth-cores into the tubes containing tracer-labelled soil made deep root nutrient uptake measurement possible up to 4.5 m soil depth. Two clear advantages of this core-labelling technique have been already addressed in detail by (Han *et al*., 2020). In short, this approach overcomes the disadvantage of the generic tracer injection, which can lead to movement of the injected aliquot depending on soil moisture content and rainfall (Hoekstra *et al*., 2014). Secondly, we were able to retract the roots of the plants that had the access to the tracers. This ensures and validates that the root activity detected in the aboveground biomass was the result of roots growing into the tracer-labelled soil of the ingrowth-cores. This eradicates the potential discrepancy between root uptake and root density data, which can potentially occur under tracer injection methods (e.g. Da Silva *et al*., 2011). However, it is unclear how well the RLD within the ingrowth-cores represent the RLD in the bulk soil around it. Our validation between aboveground tracer concentration and belowground root density showed a range of variation (R^2^ values between 0.31-0.97) leading to an average of R^2^ 0.67 (see Figure 12) confirming the reliability of the approach.

However, there was discrepancy between the two root methods, namely, RLD by ingrowth-core and pRLD by minirhizotron methods, despite the significant correlations (P≤0.005). Several factors might have led to this disagreement between the two methods. Firstly, the roots growing into the soil packed into the ingrowth-cores may not represent those growing near the inserted minirhizotron tubes well. Several soil conditions differ, and the roots in the bulk soil were typically well established before insertion of the ingrowth-cores, while the roots in the ingrowth-cores are the young roots which recently colonized the ingrowth-core soil. Also, the two-months incubation time for ingrowth-cores in this study might have been sufficient for the freshly grown fine roots inside the ingrowth-cores to undergo root turnover, whereas the roots visible to minirhizotron tubes might have been less susceptible to decomposition (Higgins *et al*., 2002), i.e., not completely exposed to soil microbes. In short, our results show that the two methods are not necessarily in agreement when determining root growth, and caution is required in interpreting the results when independently used.

### Sensor-based deep-water monitoring

The experimental setup in the DRL facility allows differentiation of crop water uptake in deep soil layers (i.e. >1 m). Perennial lupine, chicory and lucerne were found to substantially use water down to 1.5 m soil depth and even deeper water uptake (i.e. down to 2.5 m) seems to occur under perennial lupine. Such results had also been for older lucerne crops, at the same location (Clément *et al*., 2022). The slower and more shallow water use by intermediate wheatgrass and dyers woad probably stemmed from different reasons. Dyers woad is a biennial crop, it produced deep roots, but matured early in the second season, terminating its water use during June. Intermediate wheatgrass, on the other hand, is a perennial crop being more water spending. There are three drought escape strategies, (1) Escaping: usually achieved through shorter crop cycle (2) Water saving: through conservative use of water and (3) Water spending: using water and allocating more biomass to the growth of deep and abundant roots (Bodner *et al*., 2015; and references therein). Therefore, identifying crop water use strategies is of high importance to the development of cropping systems that suit environmental conditions and maximize soil moisture capture (Blum, 2009). By allowing continuous measurements of root growth, water and nutrient uptake in deep soil layers, DeepRootLab constitutes a unique facility for the study of deep root growth and functions at the field crop and agricultural crop rotation level.

### Addressing research questions

#### Root growth dynamics

Among the tested crops, only few (e.g. lucerne) have been reported for dynamics in root depth/penetration over multiple seasons. We observed an overall decrease in root penetration rate over time (see Figure 7). Nevertheless, looking at the yearly root growth dynamics (Figure 8), it was clear that these perennial crops continue to develop deeper root systems over the years. However, the rates of root depth penetration were low compared to many annual crops (e.g. Thorup-Kristensen, 2006). For many of the species studied, root depth penetration was also lower over the years of development than at the start, as shown by the intercepts of the regression lines being in the range of 0.5 to 1.5 m for most crops, except for mugwort, rosinweed and comfrey where intercepts were close to zero.

We also found a substantial variation in root penetration rate and total root depth between the species (see Figure 7). Our field site had a high water table (see Figure 13) – which might have restricted the root establishment of some of the crops, namely, lucerne, intermediate wheatgrass and perennial lupine. Indeed, a high water table has been shown to restrict root depth development of lucerne, which was limited to around 2 m of maximum root depth under unfavourable subsoil conditions (Dolling *et al*., 2005). However, other perennials, such as rosinweed, comfrey and curly dock reached a rooting depth of 3 m or more, indicating substantial species-variation in root penetration capacity under water logging subsoil condition (Jacob *et al*., 2013).

In 2018 from March to August, a severe drought with high temperatures occurred at the study site (see Figure 2). During this period, the average temperature was 18.9ºC (compared to 15.7ºC for five previous seasons) and the precipitation received was 48 mm, which was merely 23 % of the five-year average (209 mm) (Sears *et al*., 2021). Accordingly, we observed a marginal increase in deep rooting and root density for perennial lupine and mugwort and even a decrease for lucerne and curly dock from the last measurements in 2017 (Jun-Oct) to the first measurement in 2018 (May-Jun). This observation contradicts the general notion that plants tend to invest more in root growth as a response to water limitation.

However, unlike the crops in need of developing deeper roots to reach the subsoil water, our perennial crops, except comfrey, already had access to the underground water. During the time of drought in 2018 the reported groundwater levels were approximately 1-2 m from Apr to Jul (Clément *et al*., 2022; also see Figure 13E). Therefore, the reduction in root density from 2017 to 2018 might have been caused by hampered root growth (Sheaffer *et al*., 1988) or increased root turnover (Kuster *et al*., 2013) because of the drought condition in 2018.

#### Deep root-mediated nutrient uptake – crop comparison

We conducted three experiments comparing root growth and nutrient uptake potential (tracer labelling) of crop species. According to the minirhizotron imaging and analysis, perennial crops exhibited deeper rooting and higher root density than annual crops (e.g. chicory vs. winter rye in Exp 1; curly dock vs. winter wheat in Exp 3). Taproot-dominated crops (e.g. perennial lupine) grew more roots at depth, whereas fibrous root-dominated crops (e.g. intermediate wheatgrass, tall fescue and winter wheat) established more roots in shallower layers. The tendency for deeper root growth by crops with high root diameter is attributed to their capacity to overcome mechanical resistance in the subsoil layers upon penetration (Materechera *et al*., 1992). When it comes to root activity, annual crops, and relatively younger crops (chicory, dyers woad and curly dock) showed stronger capacity to grow fresh roots into ingrowth-cores leading to a greater ^15^N uptake. Older crops with higher root density at depth observed via minirhizotron tubes, tended to grow fewer roots into the ingrowth-cores, and exhibited weaker ^15^N enrichment. Rosinweed as an older crop showed greater N uptake capacity at 2.2-2.8 m.

Choice of management (e.g. early sowing, subsoil structurization) and genotypes (e.g. with high root penetration rate) can often lead to deeper and denser root systems accompanied by improved nutrient uptake potential (Han *et al*., 2015; Rasmussen & Thorup-Kristensen, 2016). It can also be driven by the differences in crop demands at the time of observation. Our data cannot address how the actual interactions between uptake capacity and demand contributed to the observed results. Nevertheless, a few points can be addressed. Firstly, annuals (winter rye and winter wheat) during the spring to summer (Mar-Jul), exhibited greater N uptake at the topsoil layer and shallower subsoil layers than did the old perennials (lucerne and rosinweed) despite their smaller root density. This is an intense growing period for newly established crops, which must assimilate N within a very short period. On the other hand, perennials can remobilize N from the previous year and have a longer season for N uptake. However, the deep-rooted perennials (e.g. rosinweed and curly dock) showed greater N uptake capacity at deeper soil layers when compared with annuals (2.2-2.8 m). This demonstrates that deep-rooted perennials have better access to deep-placed nutrients, improving overall nutrient use efficiency and hold potential for preventing nitrate leaching loss (Thorup-Kristensen, 2006; Van Tassel *et al*., 2017).

## Conclusions

We have successfully established a field facility in which a variety of measurements on deep root growth and function can be made. It allows the relatively easy and repeated study of root growth and function to great depth under field conditions, by using pre-installed minirhizotrons, access-tubes and soil water sensors. The extent and design of the facility allow experiments with up to 48 plots, allowing several species or treatments to be studied in well-replicated studies. The facility is an ideal platform for conducting studies at deep soil layers in the field with a capacity for generating statistically and biologically meaningful results.

## Data availability

The raw images, training datasets, manual counts and models generated after training are available **http://doi.org/10.5281/zenodo.15213661**.

## Author contributions

**EH** prepared the manuscript and all co-authors contributed to writing. **EH** trained the model for automatic root segmentation and conducted all the ingrowth-core experiments. **CC** installed the TDR system and conducted the experiment. **AGS** created the RootPainter software and provided technical advice on training models for root detection. **KTK** and **DBD** conceptualized the facility establishment. **KTK, DBD, EH** contributed to the experimental designs.

## Acknowledgements

This project has received financial support from the Villum Foundation (VKR023338). Upon manuscript writing EH was a Marie Curie Global Fellow working on a project SenseFuture (No. 884364) funded by European Union’s Horizon 2020 Research and Innovation Programme. EH is also a recipient of Novo Nordisk Foundation Starting Package (NNF23OC0082788) and DFF Sapre Aude Leader grant (4254-00064B). AGS is supported by Novo Nordisk Foundation grant (NNF22OC0080177). We are very grateful to Alan Hansen for co-designing and building the access-tubes and ingrowth-cores. Simone Fiil Svane has provided valuable advice on operating the minirhizotron camera. We thank the technical assistance of Jason Allen Teem, Aymeric d’Herouville and many others at the University of Copenhagen.

## Supplementary data

**Supp. Table 1.** Published studies based on the data acquired from DeepRootLab (DRL) (up to 2023)

| Source | Device used | Root depth | Tracers | Crops investigated | Treatments |
| --- | --- | --- | --- | --- | --- |
| (Han et al., 2020) | Ingrowth-cores<br>Access-tubes | 4.2 m | Li, Cs, Se, Sr, Rb | Lucerne | Soil depths<br>Location of labelling<br>Season |
| (Han et al., 2022b) | Ingrowth-cores<br>Access-tubes | 2.5 m | <sup>33</sup>P | Intermediate wheatgrass<br>Perennial lupine<br>Rosinweed<br>Curly dock | Crop species<br>Soil depths |
| (Han et al., 2022a) | Ingrowth-cores<br>Access-tubes | 4.2 m | <sup>15</sup>N, Cs | Lucerne-Winter rye<br>Intermediate wheatgrass-Perennial lupine<br>Rosinweed-Winter wheat | Intercropped vs. sole-cropped |
| (Clément et al., 2022) | TDR sensors<br>Minirhizotron tubes | 2.8 m | <sup>2</sup>H<br><sup>18</sup>O | Lucerne<br>Intermediate wheatgrass | Crop species |
| (Czaban et al., 2023) | Ingrowth-cores<br>Access-tubes<br>Minirhizotron tubes | 3 m | <sup>15</sup>N, Cs, Se | Sugar beet<br>Chicory | Intercropped vs. sole-cropped |

**Supp. Table 2.** Physical and chemical soil characteristics at the study site and for the ingrowth-cores.

| Soil type | Soil depth | pH | Clay | Silt | Fine sand | Coarse sand | Bulk density | P | K |
| --- | --- | --- | --- | --- | --- | --- | --- | --- | --- |
|  | (m) |  | (%) |  |  |  | (g cm<sup>-3</sup>) | (%) |  |
| Ingrowth-core soil |  | 8.1 | 12.5 | 12.4 | 45.5 | 28.9 |  | 0.004 | 0.041 |
| Field soil | 0-0.25 | 7.6 | 13.0 | 15.2 | 42.3 | 27.8 | 1.56 | 0.042 | 0.110 |
|  | 0.25-0.75 | 7.8 | 20.2 | 12.9 | 40.3 | 26.1 | 1.64 | 0.013 | 0.084 |
|  | 0.75-1.5 | 4.5 | 19.9 | 16.3 | 37.8 | 25.9 | 1.76 | 0.006 | 0.065 |
|  | 1.5-3.0 | 8.2 | 19.3 | 18.9 | 36.6 | 25.1 | 1.77 | <0.004 | 0.074 |
|  | 3.0-4.5 | 8.1 | 19.0 | 25.9 | 33.1 | 21.7 | 1.77 | 0.004 | 0.111 |

**Supp. Figure 1.**
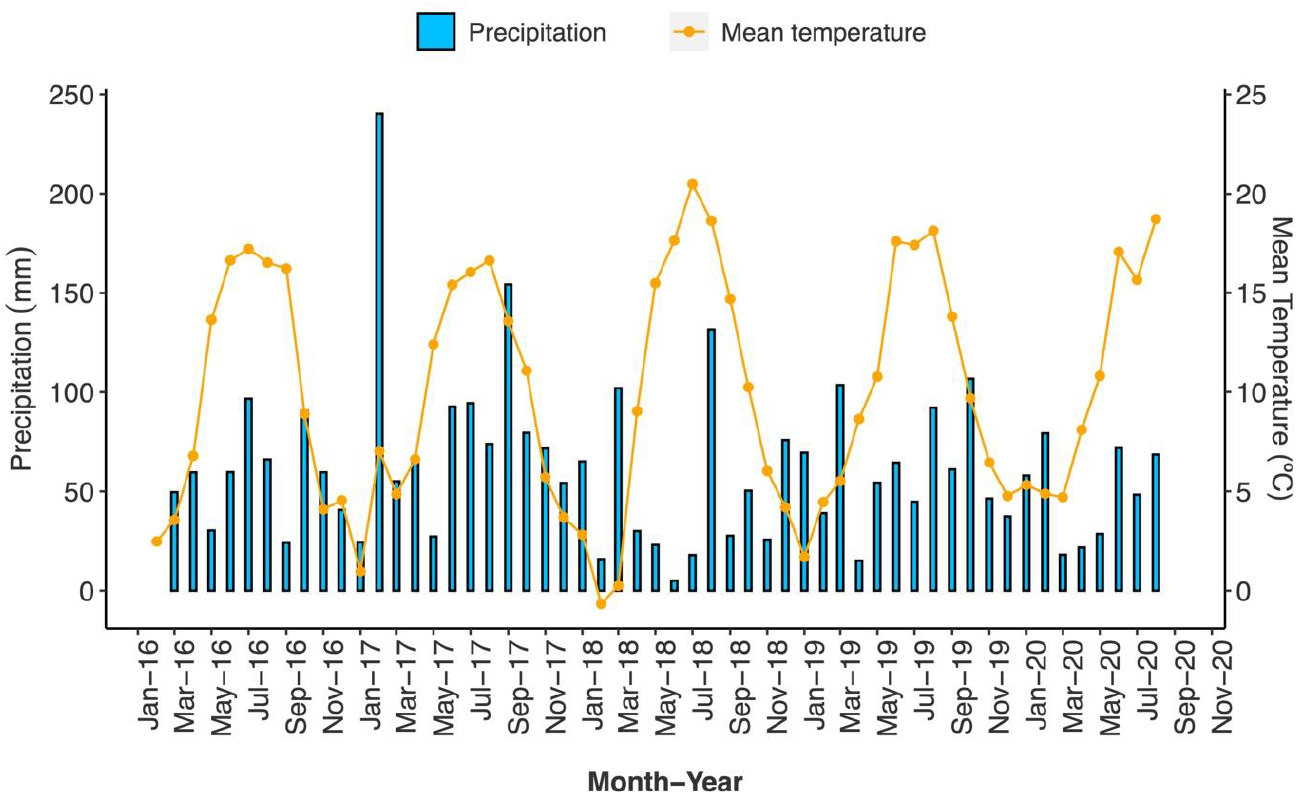
Weather conditions at the study site in Taastrup, Denmark

